# Chemically tunable permeability of engineered alpha-Hemolysin in synthetic cells

**DOI:** 10.64898/2026.05.26.727826

**Authors:** Elisabeth Bobkova, Anastasia Götz, Frank Abendroth, Olalla Vázquez, Zakarya Benayad, Viktorija Dujmović, Luis F Gutiérrez-Mondragón, Scott Scholz, Gerhard Hummer, Tobias J Erb

## Abstract

Controlling molecular transport across membranes is a defining feature of living cells, yet replicating this functionality in synthetic systems remains a major challenge. Self-inserting protein nanopores, such as α-hemolysin (αHL), offer a promising route towards programmable membrane permeability due to their robust assembly, compatibility with diverse membrane systems, and intrinsic permeability for diverse biomolecules. Here, we explored the use of chemically functionalized αHL nanopores as tunable transport modules. To quantify translocation of peptide substrates across αHL-containing membranes, we developed a high-throughput luminescence-based breakage-controlled assay using large unilamellar vesicles. With this assay we introduce a one-pot nanopore modification and strategy, compatible with scalable workflows. Electrophysiology and molecular simulations demonstrate that the introduction of cysteine residues at defined pore locations, combined with targeted chemical modification, enables controlled tuning of αHL-nanopore selectivity based on peptide structure and charge. Together, these findings position engineered protein nanopores as versatile and responsive components for controlling membrane transport in synthetic biology.

## Introduction

One of the fundamental features of living cells is their ability to precisely control molecular transport across membranes, which allows them to perform selective nutrient uptake, remove waste and sense the environment ^1^. To control molecular transport, nature evolved a complex and finely tuned system of different transporters, channels, peripheral proteins and lipids that make up the biological membrane. Despite major advances in synthetic biology, achieving the same level of control on synthetic membrane systems remains a major challenge, which limits the potential for fundamental science (e.g., construction of synthetic cells), as well as practical applications (e.g., drug delivery or bio-sensing). Current methods, such as detergent-coupled delivery of natural transporters ^2^ or the use of bacterial secretion systems for membrane protein insertion ^3,4^ allow to introduce complex protein machinery into synthetic membranes. However, they often show poor control over transporter integration and directionality, are limited to defined membrane compositions, and often interfere with other cell-free technologies, such as *in-vitro* transcription-translation (TxTl) systems ^5–9^. Engineered DNA nanopores have advanced the field by equipping membranes with size selectivity ^10,11^, and charge-based discrimination ^12^. However, their assisted assembly process, dependence on lipid anchors or chemical modification for membrane insertion, and limited chemical diversity pose similar challenges for broader implementation and application diversity ^13,14^.

An alternative strategy to above efforts is the use of self-inserting protein-based nanopores, such as alpha-Hemolysin (αHL). This approach addresses several of aforementioned limitations: nanopores assemble very efficiently and unidirectional in different types of membranes including pure block-copolymers ^15,16^, are compatible with TxTl systems ^17^, and are naturally permeable to a variety of (bio)molecules ranging from ions to nucleic acids and peptides ^18^. Furthermore, recent advances in αHL nanopore engineering demonstrated their use for membrane display ^19^, optically controlled opening and closing ^20^, while site specific cysteine mutations opened up the possibility for thiol-selective chemical functionalization, allowing for new nanoreactor functions ^21^, targeted transport between vesicles ^22^, and sensing applications ^23,24^.

Still, despite these promising advancements, self-inserting protein nanopores remain largely underexplored as a modular and scalable solution for establishing selective permeability in synthetic membrane systems. Part of this gap can be attributed to the lack of suitable assays to characterize and quantify the permeability of nanopores for different substrates. Electrophysiological assays have long been state-of-the-art, however, they are limited by their low throughput, their requirement for planar membranes, and their focus on ions ^25^. Giant unilamellar vesicle (GUV) and large unilamellar vesicle (LUV)-based assays opened the possibility to study permeability of nanopores towards biologically relevant molecules in a more realistic scenario (“artificial cell-like”) and at higher throughput ^12,25–28^. However, these assays still often require complex setups, and are challenged by the increasing instability of membranes upon pore insertion, which causes vesicle breakage that in turn causes high noise in these assays.

Here, we characterized the permeability of (modified) αHL for different substrates with a particular focus on peptide substrates. For the latter, we developed a LUV-based luminescence assay that enables direct quantification of the crossing of peptides across synthetic membranes ^29^. Using this assay, we show that introducing cysteine residues at different parts of the nanopore (i.e., pore lumen and pore entry, respectively ^21,22^) changes the permeability for peptides of different structure and charge. This selectivity can be further tuned through chemical functionalization of the cysteine residues, with molecular simulations providing insights into the mechanistic basis of selectivity. We show the controlled modification of cysteine-containing nanopores pre- and post-insertion, and present a “one-pot” modification and screening assay that is compatible with high-throughput approaches in the future. Overall, our study provides new methodological approaches and strategies into the engineering of nanopores as tunable and responsive matter for synthetic cell applications and beyond.

## Results

### Construction of alpha-Hemolysin variants and their site-specific functionalization

To enable site-specific modification of αHL, we chose to introduce cysteines into the protein and functionalize them with thiol-selective chemistries. Because native αHL does not possess any cysteines, this approach enables exquisite control over the position and timing of pore functionalization, while retaining full compatibility with in-vivo and in-vitro pore production. We designed two different αHL variants with cysteines located at two different positions: at the pore entry (0C variant), i.e., at the opening of the pore, as well as at the pore lumen (T117C variant), close to the restriction point of the pore (**Figure 1 A**). For covalent modification of the cysteine, we sought to explore two different functionalization strategies: (reversible) disulfide bond formation with reactive methanethiosulfonate reagents **(Figure 1 D)** or cysteine-peptides (**Figure 1 E**), and, on the other hand, functionalization with maleimide-based reagents, which results in irreversible bond formation between the αHL pore and the reagent **(Figure 1 F)**.

**Figure 1:**
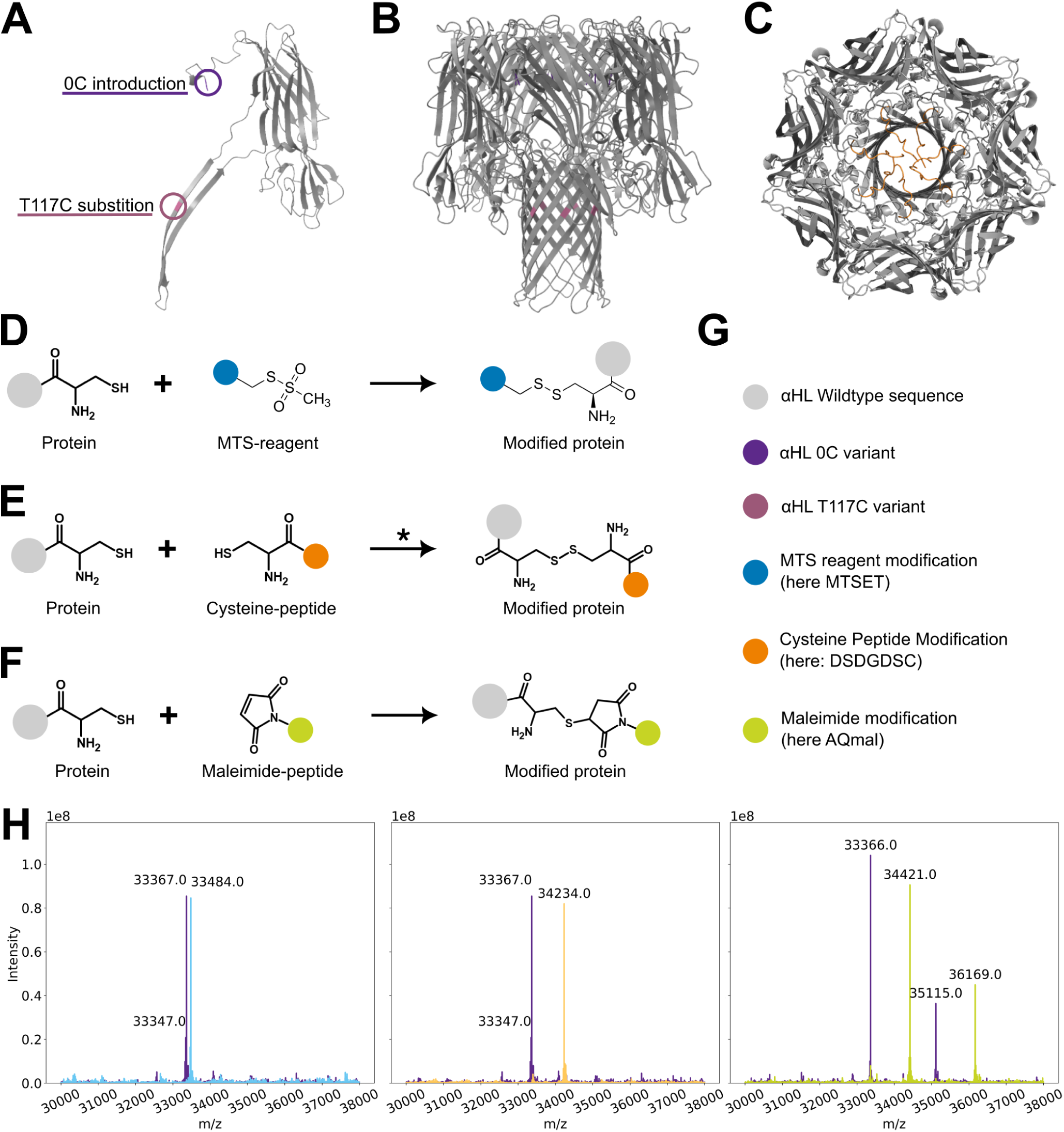
α-Hemolysin variant design, chemical modification and characterization. A) highlighted positioning of 0C (pore entry, green) and T117C (pore lumen, violet) mutations on the αHL WT monomer ^15^. B) highlighted positioning of 0C and T117C mutations on the αHL WT heptamer ^15^. C) Colabfold ^31^ prediction of a αHL heptamer with charged linkers fused to the N-terminus (highlighted in orange). In the course of this work, the mutated αHL variants were functionalized with methanethiosulfonate (MTS) reagents (D), cysteine-containing peptides (E) in presence of oxidizing agents like potassium ferrocyanide or coper(II) chloride (*), or chemically modified using maleimide-conjugated reagents (F). G) colorcodes for the *αHL* variants and modifications. H) exemplary deconvoluted masses from mass spectrometry measurements of unmodified αHL 0C variant monomers and the respective modifications: MTS-reagent in blue, cysteine-peptide in orange and maleimide reagent in green.

We prepared wild-type (WT), 0C, as well as T117C αHL monomers, incubated them with modifying reagents and catalysts, and quantified the yield of pre-insertion functionalization with intact protein mass spectrometry (MS) (**Figure 1 H**). Pre-insertion modification with cysteine-peptides (here the 8 amino acids peptide DSDGDSDC) showed different efficiencies that were depended on the type of catalyst used, ranging from ∼50% modification (CuCl_2_) to >90% (K_3_Fe(CN)_6_; PFC) (**Supplementary Figure 1 D**). Pre-insertion function-alization with sodium (2-sulfonatoethyl) methanethiosulfonate (MTSET) consistently yielded ∼100%, while functionalization with maleimide derivative AQuora® Fluorescein-Maleimide (AQmal), showed ∼85-100% efficiency depending on the pH and incubation time. Interestingly, for the pore lumen variant T117C, in few cases the reaction was performed at pH 8, we observed a minor byproduct formation with two modified sites, likely attributed to N-terminal nitrogen functionalization via aza-Michael addition ^30^ **(Supplementary Figure 1 C)**.

We also assessed post-insertion functionalization, for which we prepared a solution of large unilamellar vesicles (LUVs) that we incubated with the different αHL monomers to allow pore assembly and insertion, before adding the modification reagents. While modification with unlabeled peptides could not be reliably verified, modification with AQmal resulted in fluorescently labeled pores that could be visualized using non-reducing SDS-PAGE, demonstrating (partial) modification of the pores post insertion **(Supplementary Figure 1 E)**. Together, these experiments showed that αHL could be modified at specific positions with high yields pre-insertion (i.e., >85%) –and in case of maleimide-based modifications– also post insertion (i.e., after their incorporation into lipid vesicles).

### Pore variants show reduced ion permeability in electrophysiological measurements

To study the behavior of the different αHL variants (and their modified versions) in more detail, we used single pore electrophysiological measurements that are well-established in the field. For that, we painted planar lipid bilayer of 1,2-diphytanoyl-sn-glycerol-3-phosphocholine (DPhPC) on chips, and applied a transmembrane voltage (V_m_), followed by addition of pore monomers. Upon successful formation of a nanopore, the measured current (I) exhibits a characteristic step-wise increase, indicating movement of ions across the newly formed pore, which can be used to characterize pore behavior (**Figure 2 A**).

**Figure 2:**
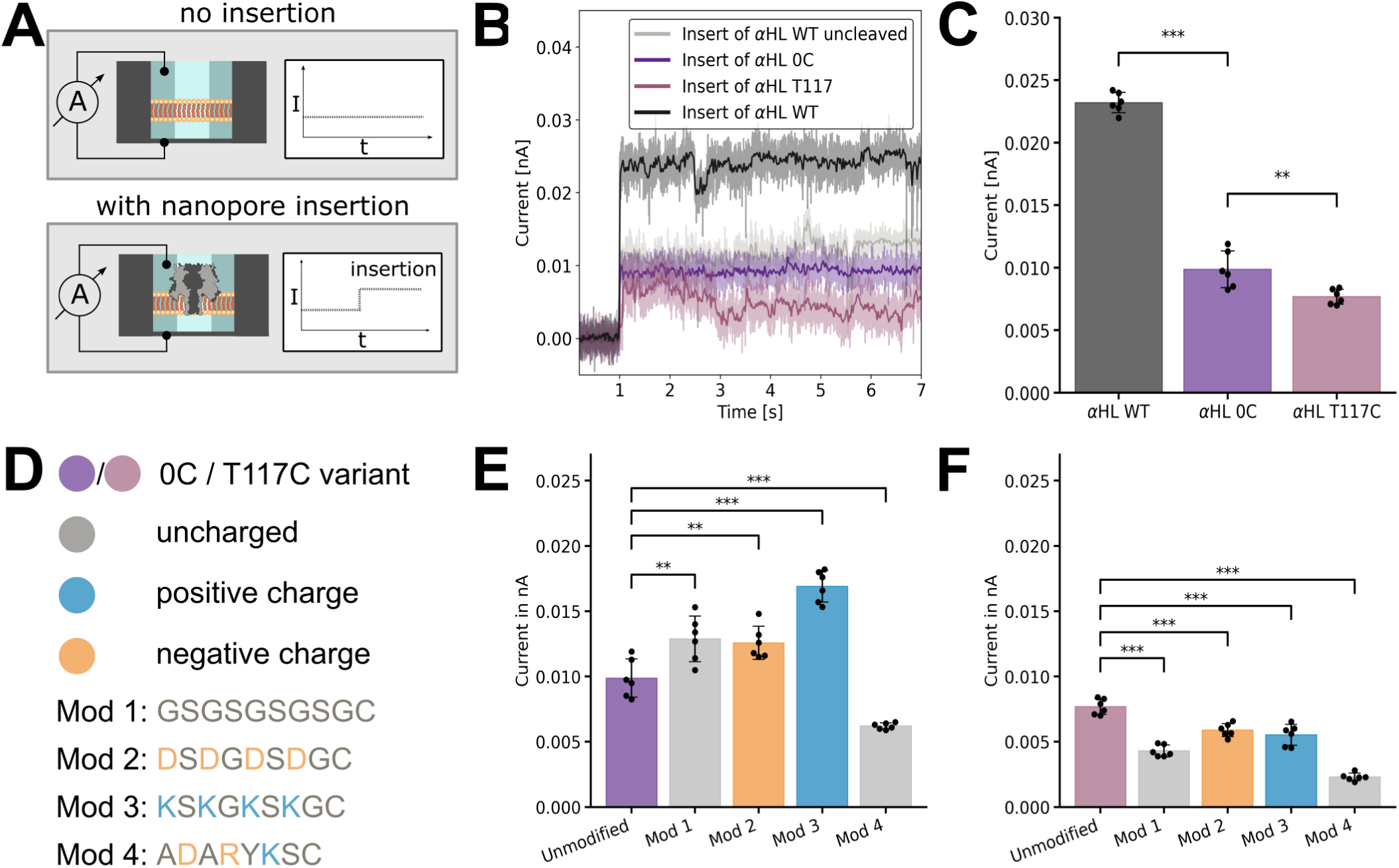
Electrophysiological measurements reveal charge and position dependent changes in the modified α-HL variant behavior. A) Schematic representation of the planar lipid bilayer current measurement setup. The planar lipid bilayer was painted over a cavity with 800mM KCl buffer allowing for current detection upon voltage application in case a nanopore insertion. B) Representative current reading for the modified and unmodified αHL entry and pore lumen cysteine variant (n=1, obtained at +30mV). C) Mean current recordings of the tested variants (n=6, obtained at +30mV). D) List of cysteine-containing peptides (mod 1-4) used for pre-insertion modification of αHL variants with the respective color-coding of charges and variants. Here, Mod 1 refers to the uncharged peptide, Mod 2 to negatively, Mod 3 to positively and Mod 4 to amphipathic peptide respectively. The amino acids carrying the charges are highlighted in blue (+) and orange (-). E) Mean current values obtained for n=6 independent insertions of the resulting modified αHL variants with cysteine at pore entry and F) with cysteine at pore lumen.

We first tested His-tagged αHL WT, as well as cleaved αHL WT, 0C and T117C variants without modification (**Figure 2 B** and **C**). The cleaved WT pore produced a stable insertion with a mean current (I_mean_ = 0.023 ± 0.0007 nA) and a current asymmetry of 1.23, which is in line with the literature ^32,33^. The presence of the N-terminal His-tag (uncleaved WT pore) led to ∼20% current decrease, which was accompanied by transient current blockages, likely through partial obstruction of the pore by the tag (I_mean_ = 0.0176 ± 0.002 nA). The mean current was further decreased in case of the 0C (I_mean_ = 0.0099 ± 0.0013 nA), and in particular, the T117C variant (I_mean_ = 0.008 ± 0.0006 nA) which is in consistent with previous studies ^34^ and in line with introduction of the cysteine residue at the restriction point of the pore. Overall, these results showed that the 0C and T117C pores were still functional, but displayed a 50 to 65% reduced ion permeability.

### Pore functionalization changes ion permeability in a position- and charge-dependent manner

Next, we modified the 0C and T117C variants with different cysteine-peptides via PFC to study the influence of pore functionalization onto ion permeability. We tested each pore variant with four different peptides that were each between 8 to 10 amino acids in length, and of neutral, positively charged, negatively charged, or amphipatic character (**Figure 2 D**). For the 0C variant, we observed significantly higher mean currents for modifications with neutral peptides (I_mean_ = 0.013 ± 0.002 nA), and negatively charged peptides (I_mean_ = 0.013 ± 0.001 nA), respectively. Introduction of a positively charged peptide, further increased the mean current to 0.017 ± 0.001 nA) suggesting that these functionalized pores were more permeable for ions compared to unmodified 0C (see **Figure 2 E**). In contrast, modification with the amphipathic peptide reduced overall conductivity of the 0C pore significantly to ∼62% of the unmodified variant (mean I_mean_ = 0.006 ± 0.002 nA), indicating that ion permeability was decreased (see **Figure 2 F**). In respect to single pore current distributions, we observed variable current broadening from standard deviations around σ = 0.0005 for uniformly charged peptides to σ = 0.003 for amphipatic peptide modification (**Supplementary Figure 2 C**).

For the T117C variant, which was functionalized at the restriction site of the lumen, all peptides independent of their charge showed a decrease of current to 30% to 77% (I_mean_ = 0.0043 ± 0.0004 nA for neutral, I_mean_ = 0.0059 ± 0.0005 nA for negative, I_mean_ = 0.0055 ± 0.0007 nA for positive and I_mean_ = 0.0023 ± 0.0003 nA for amphipatic peptide modifications). In constrast to the 0C pore, T117C showed similar current broadening at σ = 0.01 across all peptide modifications (**Supplementary Figure 2 D**).

Finally, we probed charge-dependence of pore permeability. To that end, we re-assessed behavior of the pores at different pHs, where charges were neutralized (i.e., pH 9 for positive and amphipatic peptides, and pH 10 for negative charged peptide modification respectively, **Supplementary Figure 3 B** and **C**). In all cases, charge neutralization eliminated the differences between the differently charged peptides, suggesting that pore permeability for ions was primarily dependent on the position of the modification, but could be further tuned through electrostatics via the charge of the respective peptide modification.

### Molecular dynamics simulations of cysteine peptide functionalized αHL variants

To better understand the underlying mechanisms of current changes upon cysteine peptide modifications of αHL variants, we performed molecular dynamics simulations. We first simulated αHL WT and variants 0C and T117C at different cysteine protonation states and determined the permeability for Cl^-^ and K^+^ (**Figure 3** A**, Supplementary Table 4**). Consistent with our experimental observations, the simulations showed that introducing cysteines strongly affected aHL’s ion permittivity. This was particularly pronounced for permeability of Cl^-^, which is naturally preferred by αHL over cations. For αHL WT, we computed a Cl^-^ current of 0.022 ± 0.001 nA, and reduced Cl^-^ currents of 0.020 ± 0.006 nA for protonated 0C, 0.011 ± 0.003 nA for deprotonated 0C, and 0.010 nA for T117C (±0.003 and ±0.002 nA for protonated and deprotonated cysteines, respectively), which is well in line with the experimental trend. At the same time, our simulations also showed an increase in cation preference, which partially recovered the total flux of ions to 0.027 ± 0.003 nA in protonated 0C, 0.022 ± 0.004 nA in deprotonated 0C, and 0.025 nA with a deviation of 0.003 and 0.002 nA for protonated and deprotonated state of T117C mutation, respectively, resulting in a less pronounced decrease in total current compared to our experimental data. To exclude that our simulation parameters might have caused this overestimation in total current, we computed the diffusion coefficients of ions in our simulation box, which showed that ion flux was only affected by 1 – 3% (*Supplementary Text 1*).

**Figure 3:**
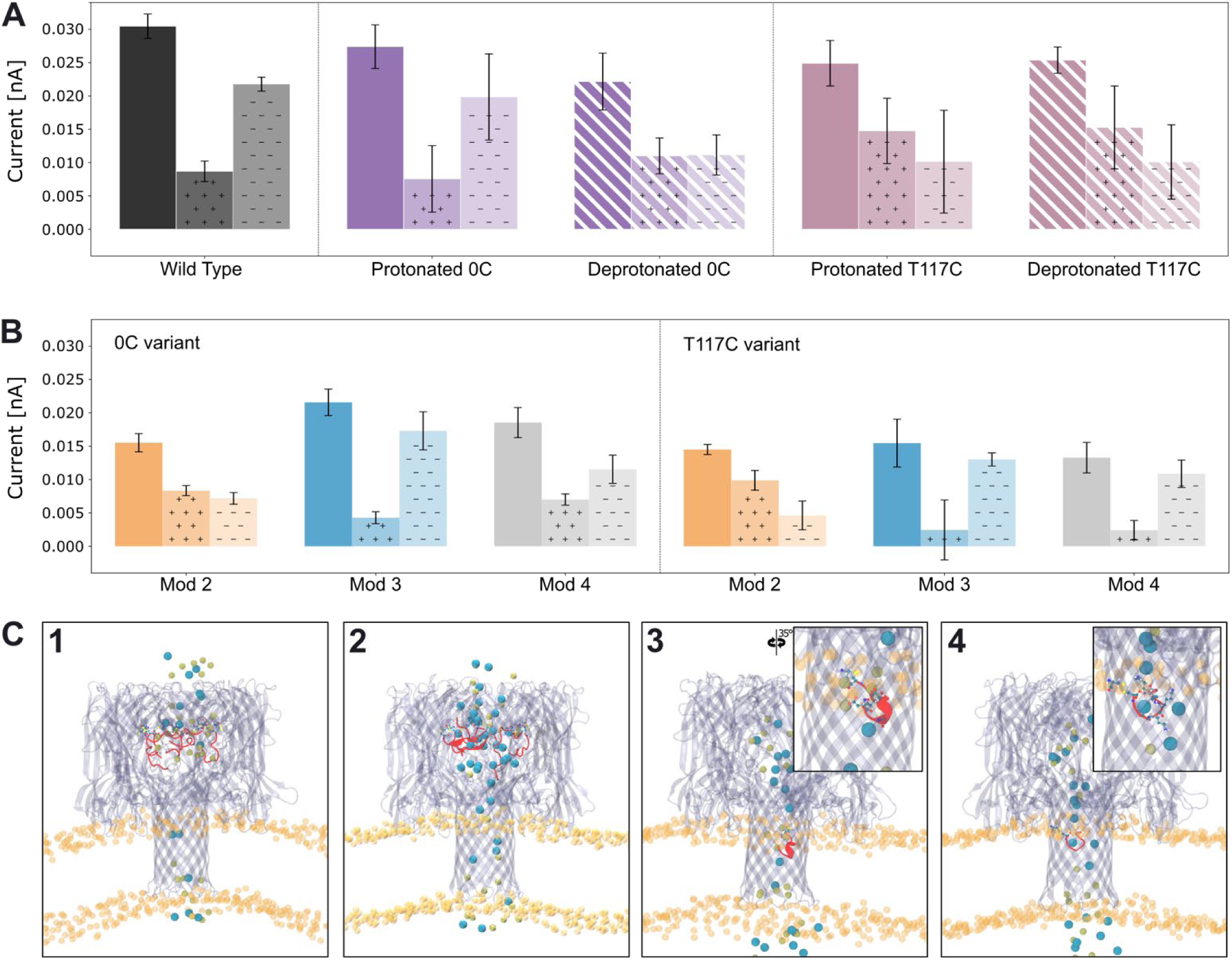
Ionic currents estimated from molecular dynamics simulations. For each system, the total ionic current is shown together with the potassium current, indicated by shaded areas with “+” symbols, and the chloride current, indicated by shaded areas with “–” symbols. (A) Comparison of WT αHL, the 0C variant with all cysteines protonated, the 0C variant with all cysteines deprotonated, the T117C variant with all cysteines protonated, and the T117C variant with all cysteines deprotonated, from left to right. Deprotonated systems are indicated by hatched shading. (B) Comparison of the peptide-functionalized systems. From left to right, systems are functionalized with the negatively charged aspartate-rich peptide DSDGDSDGC (Mod 2), shown in orange; the positively charged lysine-rich peptide KSKGKSKGC (Mod 3), shown in blue; and the amphipathic peptide ADARYKSC (Mod 4), shown in grey. (C) Snapshots from MD simulations of αHL functionalized with (1) negatively and (2) positively charged peptide in 0C, and (3) negatively and (4) positively charged peptide in T117C mutation. The protein is shown in transparent purple, peptides are shown in red, and the disulfide bond with the protein mutation is shown in CPK. Potassium ions are colored yellow, while the chloride ions are colored blue. The membrane phosphorus is shown in transparent ochre. Water is not shown for clarity. For the T117C mutation (3, 4), the zoom in on the peptide is shown, with charged residues shown in CPK.

We also computed ionic currents of variants 0C and T117C modified with amphipathic, negatively charged, and positively charged peptides. For 0C, we introduced seven peptide modifications per pore, while for T117C, only one peptide modification was introduced, as preliminary tests showed that introduction of more than one modification seemed to fully block the channel during simulation. Again, we computed the permeability for Cl^-^and K^+^, individually (**Figure 3 B, Supplementary Table 4**).

In the simulations of the 0C modifications (**Figure 3 C1,2**) accumulation of ions in the newly introduced, charged peptide mesh led to the changes in ion selectivity. For the negatively charged peptide, we observed a drop in total current (0.016 ± 0.001 nA) due to the unfavorable passage of Cl^-^ ions (0.0072 ± 0.0009 nA). However, the K^+^ current was uninterrupted at 0.0083 ± 0.0008 nA. In contrast, positively charged peptides recovered Cl^-^ selectivity of the WT channel while suppressing the K^+^ passage, leading to an increased net current of 0.022 ± 0.002 nA. Modifications with amphipathic peptides, recovered the charge profile of the WT channel, but the resulting spatial constrictions reduced total (0.019 ± 0.002 nA) and individual currents (0.0070 ± 0.0008 nA and 0.012 ± 0.002 nA for K^+^ and Cl^-^, respectively). While these simulations generally covered our experimental trends, we note that undersampled peptide conformational dynamics and changes in protonation states may further impact our results.

In case of T117C, we calculated a total current of 0.0145 ± 0.0008 nA for modification with the negatively charged peptide, 0.015 ± 0.004 nA for the positively charged peptide and 0.013 ± 0.002 nA for the amphipathic peptide. In contrast to the 0C modifications, where altered ion flux had resulted from accumulation of ions, our simulations showed that the modified charge profile of the T117C pore was caused by interactions of ions with individual amino acids on their way through the pore (*Figure 3* C 3,4). As for our calculations of the 0C modification, we expect that more extended sampling of protonation and conformational states could further improve the quantitative consistency. All in all, however, our simulations were able to cover major experimental trends—especially for the introduction of positively and negatively charged residues—and provide insights into the molecular basis of changes in ion permeability of modified pores.

**Figure 4:**
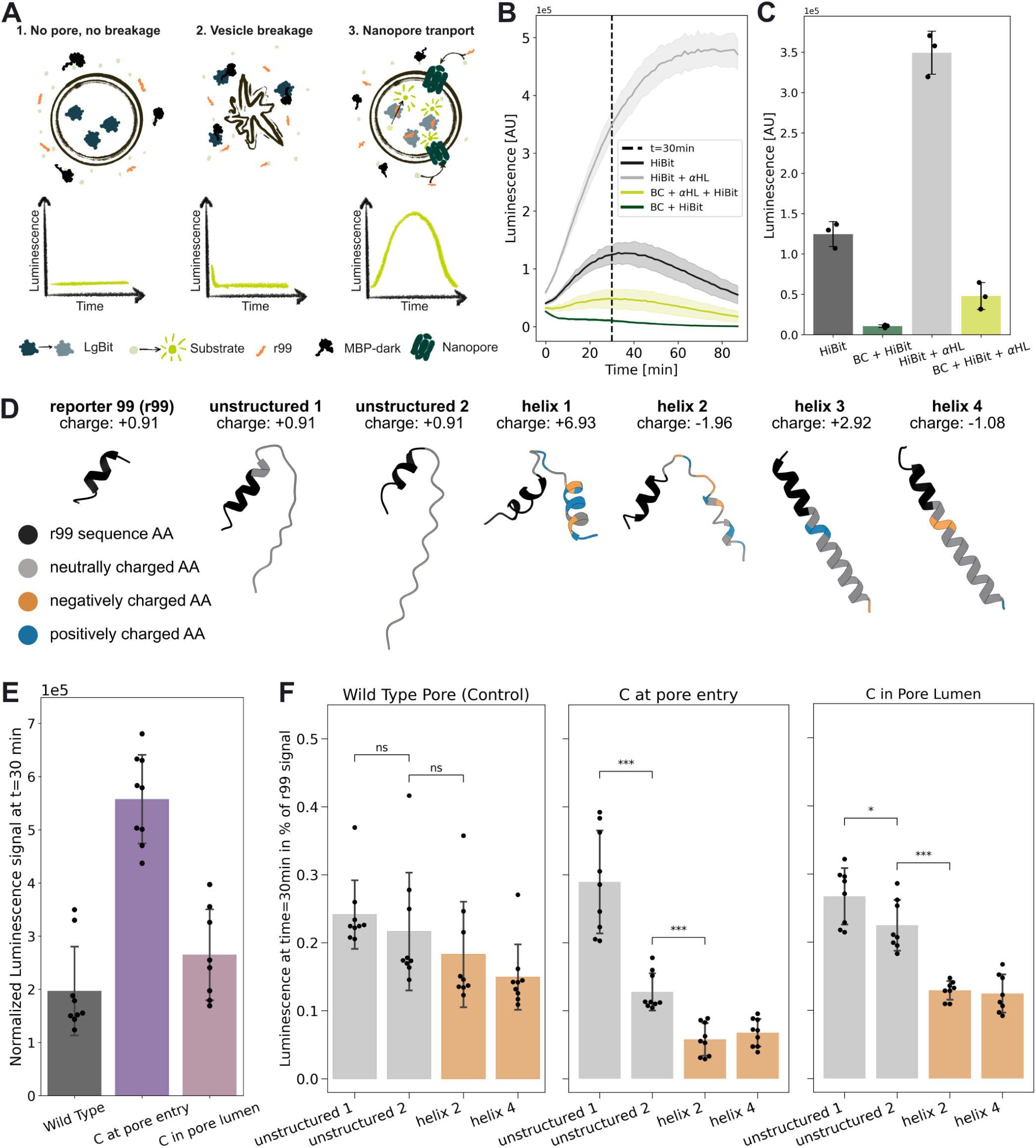
Split Nanoluc-based peptide diffusion assay reveals charge and secondary structure dependent diffusion patterns for studied αHL variants. A) Schematic representation of the breakage controlled split-nanoluc based assay for peptide diffusion analysis. In short, S11 (large bit, LgBit) is encapsulated in LUVs, while the quenching peptide fused with MBP (MBP-dark), the reporter peptide (r99) and the luciferase substrate are added to the outer solution. In case of vesicle breakage, the LgBit binds to the high affinity MBP-dark, quenching its activity. Upon addition and successful insertion of the nanopore with subsequent r99 diffusion into the vesicle, the LgBit binds r99, forming a functional nanoluc that converts furimazine to furimamide, emitting a detectable luminescence signal. B) Example luminescence output of the developed assay with (green) and without the addition of breakage control (grey) in presence of vesicles using HiBit peptide as a reporter peptide (n=3 technical replicates). C. Corresponding luminescence values at t=30min. D) Structures (generated with ColabFold ^31^) and total peptide charges of the base reporter r99, and its modified variants calculated for pH 6.5, with charged residues highlighted in orange (-) and blue (+). E) Luminescence signal recorded at t=30min and normalized to the buffer for the 3 pore variants with r99 reporter. F) Luminescence signal recorded at t=30min and normalized to the buffer and base reporter (r99) controls for tree αHL variants and 6 reporter variants (min. n=2 biological with n=3 technical replicates each, pooled).

### Pore variants show selective peptide permeability in a novel LUV-based luminescence assay

Having characterized ion permeability of (modified) pores, we next wanted to assess their permeability for other biologically relevant biomolecules, in particular peptides. For that, we sought to establish an assay that would allow us to quantify the transport of peptides across synthetic membrane vesicles, i.e., LUVs, in bulk solution.

To that end, we adopted a recently developed protein translocation assay that is based on a spilt nano-luciferase system (**Figure 4 A**) ^29,35^. The system consists of an inactive nanoluciferase 11S fragment (LgBit) that can be complemented into a functional enzyme through short reporter peptides (11 amino acids). As reporter peptides, we used the 11 amino acid peptides HiBit, and r99 that show high affinity (HiBit, K_d_ ∼1 nM) and low affinity (r99, K_d_ of ∼1 µM) to LgBit, respectively ^36^. We encapsulated the LgBit into large unilamellar vesicles (LUVs), and added the respective reporter peptide to the outer solution. Because HiBit and r99 are membrane impermeable, LgBit complementation can only take place, when the reporter crosses the LUV bilayer (e.g., assisted through a functional nanopore). This results in active nanoluciferase that is able to convert the membrane-permeable substrate furimazine into luminescent furimamide.

The encapsulation efficiency of LgBit was first tested for different LUV lipid compositions (**Supplementary Figure 4 B**), where a strong preference for the presence of negatively charged lipids was observed. Thus, all subsequent experiments were carried out using a DOPC:DOPG:DOPE (40:30:30 mol%) lipid mix. When verifying the assay with the HiBit reporter and αHL WT, we observed a strong background signal from LUV breakage and release of LgBit into the outer solution (**Figure 4 B**). To suppress this false positive signal, we employed an engineered “dark peptide” that blocks LgBit fragment activity ^35^. To prevent unintended transport of the dark peptide into LUVs, we fused it to maltose-binding protein (MBP-dark). Indeed, addition of MBP-dark resulted in twelvefold signal decrease of background activity and improved the signal to noise to fourfold, indicating that MBP-dark suppressed LgBit leaking from destabilized or broken vesicles (**Figure 4 B** and **C**).

With the optimized assay established, we next evaluated the r99 reporter peptide using the WT, 0C, and T117C αHL pore variants. All pores produced a luminescent signal, indicating successful translocation of r99 across the LUV bilayer, with 0C variants showing the highest apparent translocation rate (**Figure 4 E**). The differences between the variants are likely originating through both, the active protein concentration variability and the reporter diffusion differences due to cysteine addition to the pore.

Next, we created different reporter peptide variants by fusing r99 to six differently charged peptides containing various secondary structure elements – from unstructured, disordered to α-helical (**Figure 4 D**). We first confirmed that the fusion proteins were still able to complement LgBit in bulk solution (**Supplementary Figure 4 C**), before testing their transport across the bilayer through the WT, 0C, and T117C αHL pores. The reporter signal was further tested in bulk in presence of LUVs (**Supplementary Figure 4 D**), where a significant drop in signal was observed for all positively charged peptides, suggesting interference of the negatively charged lipids with the assay. Thus further analysis was performed only with neutral and negatively charged reporter variants. To account for the various pore variant and batch-to-batch differences, we normalized the luminescence signal of the reporter peptides against the r99 signal for each pore variant individually (**Figure 4 F**).

All tested reporter variants were able to cross the bilayer in the presence of the different αHL pore variants (i.e., WT, 0C, or T117C). Especially long unstructured and uncharged reporters crossed the membrane with >20% relative to the r99 reporter only signal acquired for each pore separately, independent of the pore variant tested, with the exception of the proline reach unstructured reporter for the cysteine at pore entry variant (∼13% relative to r99 reporter). Negatively charged reporter peptides passed bilayers at almost comparable efficiency in case of the WT (∼18%), while membrane crossing for these peptides was strongly reduced in case of the T117C (∼12%) and 0C pore variant (∼7%). These findings are in line with earlier reports that showed that the WT and αHL variants are permeable to neutral and negatively charged peptides of different structures ^37,38^. At the same time, our results demonstrated that introduction of specific mutations can further influence the selective permeability of pores for different peptides.

### Functionalization tunes selective peptide-permeability of pores

Having shown that the different pore variants possessed some (intrinsic) selectivity for different peptides, we finally wanted to test whether this selectivity could be further tuned through pore functionalization. We were especially interested in post-insertion modifications, because this would allow temporal control over modification and minimize any differences in pore insertion and assembly compared to the unmodified pore. We first assessed the influence of different functionalization chemicals onto the nanoluciferase assay (**Supplementary Figure 4 A**). Because MTS and PFC reagents strongly suppressed luminescence signals, we decided to use maleimide-based post insertion pore functionalization with AQ-mal (**Supplementary Figure 4 A**), which lowered signal intensity only about twofold.

We slightly adopted our LUV-luminescence assay by adding a two hours AQ-mal functionalization step after pore insertion, before addition of the different reporter peptides and luciferase substrate (**Figure 5 A**). Again, we used normalization with the r99 reporter peptide to correct for individual differences between pore variants and/or any batch variations (see above).

**Figure 5:**
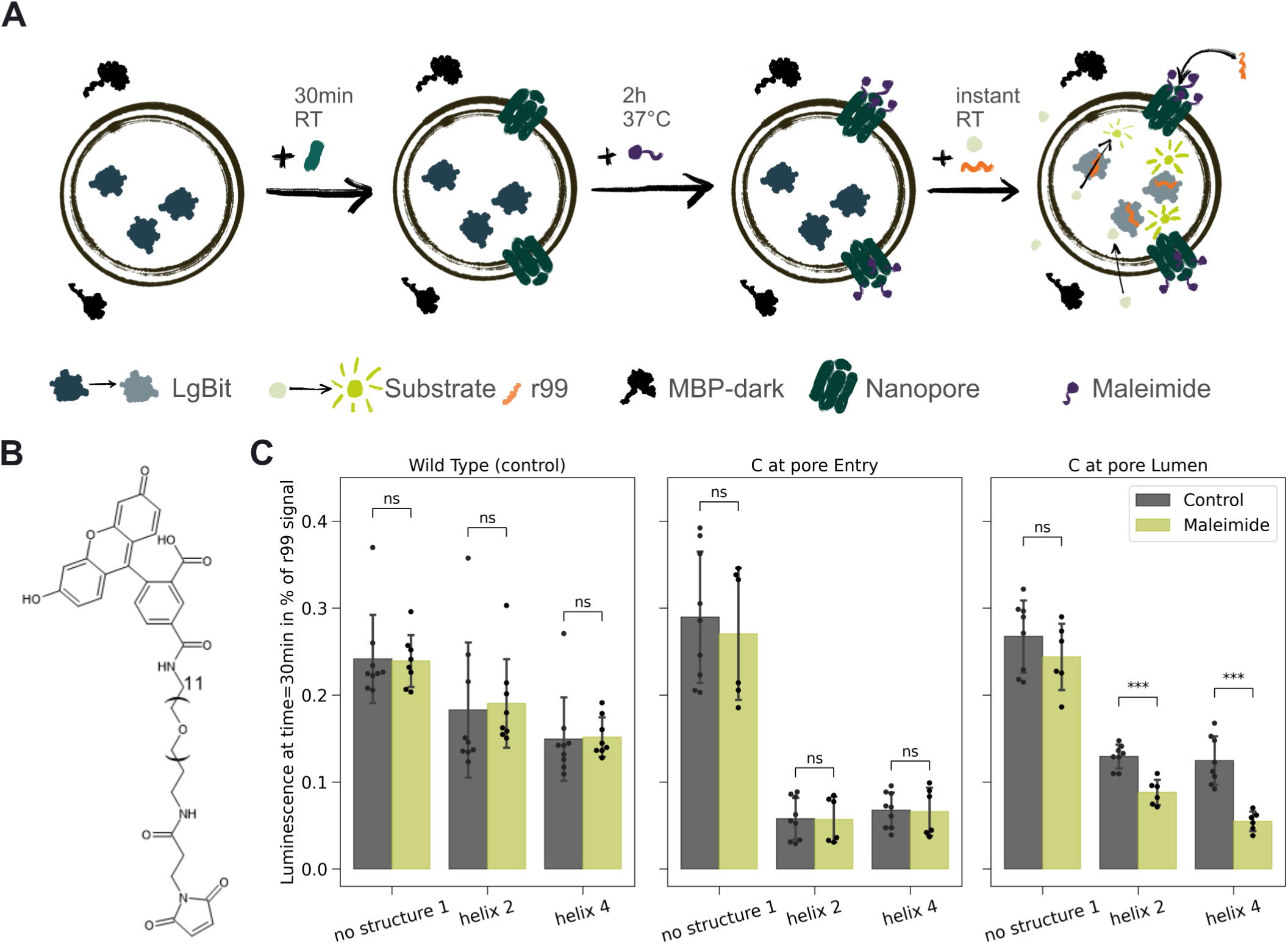
AQmal addition changes the permeability of alpha-Hemolysin variants in a position dependent manner. A) Nanopore modification combined with peptide diffusion workflow: reporter-vesicles are first incubated in presence of αHL to facilitate insertion, with subsequent modification with maleimide post-insertion for 2h at 37°C and later addition of the reporter peptides with the luciferase substrate directly before measurement start. B) Structure of AQmal employed for the variant modification (structure created using ChemDraw) C) Mean luminescence output of the maleimide-conjugate treated (green) and untreated (grey) αHL variants, taken at t=30min and normalized to the background and reporter sample signal as described in the methods section (data pooled from minimum n=2 independent assays with n=3 technical replica).

Notably, AQmal functionalization of the WT and 0C pores had no significant influence on the permeability patterns for neutral, unstructured and negatively charged peptides (**Figure 5 B**). However, functionalization of the T117C pore specifically reduced permeability for negatively charged reporters up to twofold (helix 4), without affecting the permeability pattern for the uncharged unstructured reporter signal. Together, these results demonstrated that nanopore permeability can be tuned in a functionalization-dependent and position-sensitive manner, validating the feasibility of this approach for engineering tunable selective diffusion of biomolecules across synthetic lipid membranes.

## Discussion

The rational engineering of selective molecular transport across synthetic membranes remains a critical challenge in biotechnology. Here, we developed new approaches to show that αHL nanopores possess some selectivity peptides (and ions) that can be further tuned through pore modification and functionalization. To that end, we established a LUV-luminescence based assay that allows to directly quantify peptide diffusion into LUVs across self-inserting nanopores. Critically, our assay is compatible with pre- and post-insertion functionalization of pores and provides an integrated breakage control, which decouples LUV breakage from true diffusion signal. Thus, it minimizes any effects from LUV instabilities or active protein concentration differences – a critical advancement over previous methods. In line with previous reports ^27^, our data confirm substantial breakage of LUVs in presence of nanopores, underscoring the need of controlling and correcting for membrane integrity in nanopore studies.

Our results validate that αHL is permeable for unstructured and α-helical peptides, but selective for neutral and negatively charges ^37^. At the same time, we provide new insights by demonstrating that the permeability for negatively charged peptides can be influenced through introduction of cysteine residues and further tuned through functionalization of the nanopores with maleimide reagents, demonstrating the potential for rational design of differential diffusion through modified nanopores. These changes are supported by molecular dynamics simulations, which provide a molecular mechanistic view of ion permeation through functionalized nanopores, and can help in further design efforts.

Overall, our work opens up new directions for engineering permeability across self-inserting nanopores. The modularity of our assay allows for high-throughput screening of (modified) αHL pore variants, and can be further extended to other functionalization strategies (e.g., disulfide bond catalysis via protein disulfide isomerase ^39^), as well as other pores, such as γ-hemolysin ^40^. This could set the stage for achieving programmable selective diffusion of individual molecules across synthetic membranes in the future.

## Methods

### Chemicals and Reagents

The relevant genes were synthetized TWIST bioscience, primers by Eurofins and peptides were produced in-house using solid-state synthesis. Lipids were purchased as chloroform solutions (Avanti). Q5 High Fidelity DNA Polymerase 2x Master Mix, NEBuilder HiFi DNA Assembly Master Mix, DpnI and chemically competent NEB Turbo and BL21 DE3 *E. coli* were purchased from New England Biolabs. NucleoSpin Plasmid mini prep and PCR cleanup kits were supplied by Macherey-Nagel. SDS-lane non-reducing sample buffer (x5), ReadyBlue^TM^ staining solution and Mini-PROTEAN^R^ TGX^TM^ precast gels were purchased from Bio-Rad Laboratories, while the Pierce^TM^ BCA-assay and Spectra^TM^ multicolor broad range protein ladder were obtained from Thermo Fisher Scientific. The Nano-Glo HiBit Extracellular Detection System with the HiBit substrate was acquired from Promega and AQuora® Fluorescein-Maleimide (AQmal) was purchased from Merck. (2-(Trimethylammonium)ethyl) Methanethiosulfonate Chloride (MTSET) and Sodium (2-Sulfonatoethyl)methanethiosulfonate (MTSES) were obtained from Biomol. Further chemicals used in this study were purchased from Carl Roth Chemicals unless indicated otherwise.

### Cloning, Gene expression and Protein preparation

αHL WT was cloned into the pET51b and combined with a T7 promotor and terminator to avoid cytotoxicity. The 0C and T117C mutations were produced via Gibson assembly and successful clones verified via full plasmid sequencing (relevant sequences are available in supplementary materials, and plasmids available on EDMOND).

αHL variants as well as MBP-dark and 11S (LgBit) proteins were purified via N/C-terminal His-tags. In brief, the respective plasmids were transformed into chemically competent BL21 DE3 *E. coli* cells (NEB) and colonies were grown overnight at 37°C on lysogeny broth (LB) agar plates supplemented with the respective antibiotic (100µg/ml ampicillin, 50µg/ml kanamycin). Subsequently 2L of terrific broth (TB) supplemented with the respective antibiotic were directly inoculated with a picked colony from the plate and grown overnight at 37°C at 120rpm shaking. The culture was induced with 500µg/ml IPTG and 0.125% w/v arabinose (VWR Life Science) and incubated for another 4h at 30°C and 120rpm. The cells were harvested at 4°C 4500g and the pellet resuspended in double volume (relative to weight) of Buffer A (50mM TRIS pH 8, 500mM KCl, 2mM MgCl_2_, 1% v/v TRITON-X100, 1x recombinant DNAseI) via vortexing. The cells were lysed via sonication at 30-50% amplitude 3 times for 1 min or until color and viscosity change (1s on, 1s off protocol; Sonoplus GM200, BANDELIN equipped with a KE76 tip). The resulting lysate was then centrifuged at 25000g for 30min at 4°C and the pellet discarded. The supernatant was filtered using a 0.45µm syringe filter (Sarstedt) before loading on a Ni-NTA agarose bead packed gravity flow column (Protino, Machery-Nagel) pre-equilibrated in buffer B. The beads were then washed 5x bed volume with buffers B (50mM TRIS pH 8, 500mM KCl) and C (50mM TRIS pH 8, 500mM KCl, 50mM Imidazole) to elute unspecific binders. The target protein was then eluted in 2x bed volume of buffer D (50mM TRIS pH 8, 500mM KCl, 500mM Imidazole). The proteins were concentrated using amicon spin concentrators (15 to 30 kDa MWCO depending on the protein size, Sigma-Aldrich), desalted using PD-10 desalting columns (Cytiva), eluted into buffer E (TRIS 20mM pH 8, 10% glycerol), snap frozen in liquid nitrogen and stored at −70°C until further use. All steps after centrifugation were performed at room temperature (RT).

### Single pore electrophysiological measurements

A 10mg/ml solution of DPhPC lipid was prepared via aliquoting the chloroform solution in a glass vial, drying for 2h under vacuum, resuspending the resulting film in n-Octane (Sigma Aldrich) and sonicating for 30min in a water bath at RT and subsequently heating the solution at 65°C for 10min. The measurements were performed on the NanIon Orbit Mini device (Nanion Technologies), on a MECA4 chip with 100nm cavity diameter at 25°C (set using Nanion temperature control unit). The cis-chamber was filled with 150µl of recording buffer (800mM KCl 100mM HEPES pH 7.5). During pH change measurements HEPES buffer was used for pH 7.5 and 8, while Glycine was used for pH 9 and CHES buffer was used for pH 10. A lipid bilayer was formed by painting the prepared DPhPC lipid solution on top of the cavity using a pipette tip, with bilayers of 10 to 45pF resistance used for the experiments. After formation, the base bilayer current was measured at +30mV and 1.25kHz sampling rate, with 1µl of 7µg/ml protein solution added to the chip chamber later. The protein insertion was promoted with single pulses at +100mV. After successful protein insertion the buffer solution was exchanged 3-5 times to prevent multiple insertions. Prior to analysis current measured for each insertion was normalized to the baseline by subtracting the mean current over 1s before insertion.

For IV recordings, the preinstalled voltage program 3 (EDR software) was used with a range from −50mV to +50mV in 5 steps of 10mV (Settings: V_hold_ = 0mV, V_pulse_ = 50mV, V_step_ = 10mV, T_hold_ = 500ms, T_pulse_ = 200ms, N = 5). Due to the limited stability of planar lipid bilayers and the oxidative sensitivity of our setup, this analysis was only carried out for αHL pore variants modified pre-insertion.

### System preparation and molecular dynamics simulation settings

The mutations and initial peptide placements were generated using the AlphaFold 3 server ^41^. For the T117C mutation in the pore lumen, only one peptide was introduced, as preliminary tests showed that adding more peptides blocked the pore and abolished the ionic current. For the 0C mutation, at the pore entry, seven peptides could be introduced while maintaining ionic flow. The resulting models were visually inspected and compared with the wild-type αHL structure, PDB ID 7AHL ^42^, to confirm good overlap of the overall protein fold. The systems were then prepared with CHARMM-GUI ^43^ using a lipid membrane composed of 100% PhPC, with a potassium chloride concentration of 0.8 mol/L. Residue protonation states were assigned for pH 7, unless stated otherwise. Simulations were performed using the CHARMM36m force field ^44^, and GROMACS 2025.1 ^45^ software. Energy minimization was performed using the steepest-descent algorithm for up to 5000 steps, with positional restraints applied to the protein backbone, side chains, lipids, and dihedral angles. Each system was subsequently equilibrated through six restrained MD stages, during which the positional and dihedral restraints were gradually reduced. The first three equilibration stages used a 1 fs time step for 125 ps each, followed by three additional stages using a 2 fs time step for 500 ps each. Temperature was maintained at 298.15 K using the velocity-rescale thermostat ^46^, with separate coupling groups for solute, membrane, and solvent. From the third equilibration step onward, semi-isotropic pressure coupling was applied at 1 bar using the C-rescale barostat ^47^. Long-range electrostatics were treated with PME ^48^, with a 1.2 nm real-space cutoff, and van der Waals interactions were treated using a force-switching scheme between 1.0 and 1.2 nm. Bonds involving hydrogen atoms were constrained using the LINCS algorithm ^49^. After equilibration, the simulation boxes were approximately 14 × 14 × 16 nm³ for the different systems.

For the systems containing the peptides DSDGDSDGC and ADARYKSC, positioned either at the pore entry or in the pore lumen, and KSKGKSKGC positioned at the pore entry, an additional 1 μs NPT equilibration was performed using a 2 fs time step under the same temperature and pressure coupling conditions. During this long equilibration, an external electric field was applied along the membrane normal using the GROMACS electric-field implementation ^45^, corresponding to an applied transmembrane voltage of approximately 30 mV. Production simulations for these systems were then performed in the NVT ensemble, starting from the final frame of the 1 μs NPT equilibration. The NVT production simulations used the velocity-rescale thermostat at 298.15 K, the same nonbonded interaction settings, LINCS constraints, and electric field applied along the membrane normal. Analysis was performed over three independent NVT replicates of 1 μs each. For the mutated αHL systems without peptide, and for the systems containing the KSKSKSGC peptide, no additional NVT production simulations were performed; instead, the 1 μs NPT simulations with the applied electric field were used directly for analysis, excluding the first 200 ns as equilibration.

#### Computation of the ionic current

The ionic current was computed using the protocol of Aksimentiev *et al* ^50^. The instantaneous current for a given ion species was computed as:

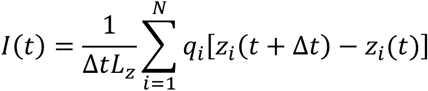

where *q*_*i*_ is the charge of ion *i*, *z*_*i*_ its coordinate along the membrane normal, *N* is the number of considered ions, and *L*_*z*_(*t*) is the instantaneous box length in the *z* direction. A time interval Δ*t* = 10 ps was used ^50^. Potassium and chloride ions were computed separately, and the total ionic current was obtained by summing the potassium and chloride contributions. This allowed us to compute the contribution of each ion type to the total current, and hence estimate the ion selectivity of the pore. The instantaneous current was integrated over time to obtain a cumulative current curve, and the average current was determined from the slope of a linear fit to the cumulative curve (**Supplementary Figure 5**). For the systems containing the peptides DSDGDSDGC and ADARYKSC, positioned either at the pore entry or in the pore lumen, and KSKGKSKGC positioned at the pore entry, the values reported in Figure 1 correspond to the mean and standard deviation of the average current calculated over three independent 1 μs NVT trajectories. For the mutated αHL system without peptide, and for KSKGKSKGC positioned in the pore lumen, the average current and standard deviation were obtained by block averaging the last 800 ns of the NPT simulation, using four blocks of 200 ns.

### Protein cleavage and modification

The αHL variants were cleaved using TEV protease (NEB) for 1.5h at 30°C. The solution was cleared of the linkers, protease and uncleaved protein by mixing NiNTA-beads in 20mM TRIS pH8 buffer solution (1:1), incubating for 5min at RT, spinning down using a table centrifuge and subsequently transferring the supernatant to a fresh vial for further processing.

For peptide modifications, 30µM protein was mixed with a 1/10 dilution of the peptide stock 10mM, to a final concentration of 1000µM or roughly 30x molar excess of the respective peptide. Then, for the cysteine-containing peptides, additionally Potassium Ferrocyanide (PFC) was added to a final concentration of 100µM to promote disulfide bond formation, while the maleimide-containing reagents did not require any further additions. The resulting protein mixes were incubated at 37°C for a minimum of 30min for cysteine peptides and 2h for the maleimide modification. For single pore current measurements, the excess reagents were removed using PD MiniTrap^TM^ G-25 columns (Cytiva) according to the manufacturer’s protocol.

### Reporter system benchmarking

In all bulk luminescence measurements the LgBit solution was initially mixed with the respective reagent, incubated and then mixed with the reporter peptide and luciferase substrate directly before the start of the measurement. For AQmal tests, a 0.78µM LgBit solution was combined with 0.03mM AQmal. A reporter solution of 20nM r99-peptide and 1/31.5 luciferase substrate dilution was mixed. The protein mix (4.8µl per well) was then combined with the substrate-peptide mix (1.2µl per well) in a white 384 well-plate (#784904, Greiner) and the plate reader measurement was directly started (Infinite 200 Pro, Tecan, Luminescence protocol at 100ms, measurement every 1.5min for 1h). For metal sensitivity measurements 7.8µM LgBit solution was mixed with 1mM of the respective metal salt, incubated at RT for 30min and subsequently combined with the peptide-substrate mix to a final concentration of 2µM of peptide and 1/125 commercial HiBit substrate (Promega) dilution. 6µl of this solution were placed in the well-plate and measured as described before. All samples were measured in technical triplicates on one plate to account for pipetting errors.

### Peptide diffusion assay

The production of LUVs for the peptide diffusion assay was based on the protocol published by Meier et al.^29^. In brief, DOPC:DOPG:DOPE lipids were mixed to a 40:30:30 molar ratio in a glass vial and dried for at least 1h in a desiccator. The dried lipid film was rehydrated with the inner aqueous (IA) solution (20mM TRIS pH 6.5, 200mM Sucrose, 3% w/v Glycerol and 15.6µM purified 11S protein (LgBit) stock) to a final lipid concentration of 10mg/ml at RT for 30min with occasional gentle vortexing. After this, the solution was freeze-thawed 5 times alternating between liquid nitrogen and a water bath at room temperature to increase protein encapsulation. In parallel, the mini-extruder (Avanti Research) was assembled according to the manufacturer’s’ protocol with a 100nm diameter polycarbonate membrane (GE Healthcare) and equilibrated with the osmolarity matched outer solution (here - 5% w/v glycerol in 20mM TRIS pH 6.5). The liposome solution was extruded 13 times and washed 3 times by centrifuging the sample for 1h at 4°C, an average of 112644 rcf, and resuspending the pellet in the OA to remove the unencapsulated protein. The final LUV pellet was subsequently gently resuspended in 400µl of OA. The resulting mix was stored in the fridge at 4°C and used within 3 days. The empty LUVs were produced using the same protocol, but here LgBit was omitted from the IA solution.

For the peptide diffusion assay, a nanoluc-based real-time kinetic translocation assay ^29^ was adapted for the use with self-inserting nanopores. Briefly, the liposomes contain the catalytically inactive 11S (LgBit) variant of the nanoluc. The split-nanoluc is completed through 11S binding the medium-affinity complementation tag (pep99) ^36^ upon its successful translocation into the liposome lumen. Its unlocked catalytic activity leads to the conversion of the substrate leading to the emission of a detectable luminescent signal. A collection of structurally distinct peptides fused to r99 was produced via solid state synthesis and used to compare the influence of charge and structure on the translocation. To selectively eliminate the liposome breakage signal, the maltose binding protein (42kDa) was fused with the high affinity 11S inactivating peptide ^35^ (plasmid kindly provided by Markus Meier) and combined with the OA solution.

For a single sample for the peptide diffusion assay, 5µl of the LUVs solution were combined with 5µl of purified MBP-dark, 8µl outer solution buffer and 1µl of αHL or buffer sample. The solution was gently mixed and left at RT for 30min to complete the self-insertion of the nanopore. Following that the maleimide reagent of choice was added to a final concentration of 22.9µM, the solution was gently mixed and incubated at 37°C for 2h. During this time the reporter-peptides were separately mixed with the HiBit substrate solution to a final concentration of 20nM peptides and 1/31.5 substrate dilution. The LUV-protein mix (4.8µl per well) was then combined with the substrate-peptide mix (1.2µl per well) in a white 384 well-plate (#784904, Greiner) and the plate reader measurement was directly started (Infinite 200 Pro, Tecan, Luminescence protocol at 100ms, measurement every 1.5min for 1h). All samples were measured in technical triplicates on one plate to account for pipetting errors. All experiments were performed at least 3 times independently.

### Statistical analysis

All data analysis was performed using Python. All data presented are the mean of *n* = 3 technical replicates ± SD, unless otherwise stated in the respective caption.

Electrophysiological data were analyzed to quantify single-channel currents following protein insertion. Pore insertion events were identified manually, and the resulting current signal was normalized to the pre-insertion baseline to correct for drift and background noise. Specifically, the normalized current, I_norm_(t), was calculated as:

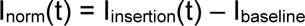

where I_insertion_(t) represents the current at time t following pore insertion, and I_baseline_ is the mean current measured during a 1-second interval preceding insertion. After that the current was filtered using Gaussian filter (σ=10, pyabf library). The mean single-channel current, I_mean_, was then calculated from the normalized current over a 5-second post-insertion window, and the standard deviation, σ, was computed over the same interval to quantify current fluctuations.

IV curves were generated from a specific voltage range by categorizing the corresponding current values for each applied voltage, ensuring that each voltage level had its own set of current data. For each dataset, the average current and its standard deviation were determined. A linear regression analysis was subsequently performed using the average current values derived for each voltage. For this analysis, three traces (N = 3) were recorded for each pore variant, and the mean IV curve was calculated from these traces. Before conducting the analysis, the data underwent processing with a Gaussian filter to minimize noise, and all IV curves were normalized the baseline signal prior to pore insertion. Each I-V curve was recorded over time spans of at least 30 seconds.

For peptide diffusion assays, data normalization was employed to account for variations in experimental conditions and reporter peptide efficiency. The normalized luminescence signal, L_norm_(t)_i_ for each technical replica i, was calculated as:

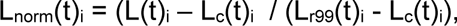

where L(t)_i_ represents the luminescence signal at time t for technical replicate i following peptide addition, L_c_(t)_i_ is the luminescence signal in the absence of peptide (control), and L_r99_(t)_i_ is the luminescence signal obtained with the r99 reporter peptide, serving as a maximum signal reference. This normalization procedure effectively corrects for background luminescence and variations in reporter peptide activity, enabling quantitative comparison of peptide diffusion rates.

## Acknowledgements

We would like to thank Markus Meier, Dr. Nataliya Safronova, Dr. Beau Dronsella, Dr. Sebastian Barthel, Dr. Owen Jarman, Dr. Martine Ballinger, Max Hoffmann-Becking, Dr. Max Gantz, Nitin Bohra, Tobias Krug, and Dr. Victoria Sajtovich for regular discussions, ideas and feedback on the manuscript. We also thank Dr. Kerstin Göpfrich, Dr. Victor Sourjik and Dr. Jan Goldschmidt for feedback.

## Author Contributions

All authors contributed to the design, execution, interpretation of the experiments, and writing and revision of the manuscript.

**EB, SAS** and **TJE** conceived the work.

**EB, ZB, VD, SAS** and **TJE** wrote the manuscript.

**EB** led the experimental work, including the adaptation of the luminescence assay, nanopore purification, functionalization, and characterization, diffusion assays, data analysis, data visualization, and experimental design.

**AG** performed the electrophsiological experiements and statistical analysis of the corresponding results.

**VD** and **ZB** performed MD simulations and the corresponding statistical analysis.

**LFGM** contributed to nanopore functionalization and characterization experiments.

**FA** and **OV** performed peptide synthesis.

**SAS** participated in experimental design.

Funding acquisition: **SAS, GH, TJE**

All authors contributed to the writing and approval of the manuscript.

## Data availability

All data are available in the main text or the supplementary materials. LgBit plasmid data has been previously published by Viereckt et al. ^51^, while other key plasmid sequences used in this study are available on EDMOND (https://doi.org/doi:10.17617/3.3U1KSM).

