## Supplementary Information for "Chemically tunable permeability of engineered alpha-Hemolysin in synthetic cells"

### Supplementary Data for the Article "Chemically tunable permeability of engineered alpha-Hemolysin in synthetic cells"

*Corresponding author:

#### Supplementary tables

**Supplementary Table 1: αHL variant sequences used in this study.** The linker is shown only for the WT sequence, where the colored part is removed after TEV cleavage (orange denotes the 6xHIs tag, blue highlights the TEV cleavage site). The 0C and T117C are shown post-cleavage, and the cysteines are highlighted in violet.

| Variant | Sequence |
|---|---|
| **WT** | MHHHHHHENLYFQSDSDINIKTGTTDIGSNTTVKTGDLVTYDKENGMHKKVFYSFIDDKNHNKKLLVIRTKGTIAGQYRVYSEEGANKSGLAWPSAFKVQLQLPDNEVAQISDYYPRNSIDTKEYMSTLTYGFNGNVTGDDTGKIGGLIGANVSIGHTLKYVQPDFKTILESPTDKKVGWKVIFNNMVNQNWGPYDRDSWNPVYGNQLFMKTRNGSMKAADNFLDPNKASSLLSSGFSPDFATVITMDRKASKQQTNIDVIYERVRDDYQLHWTSTNWKGTNTKDKWTDRSSERYKIDWEKEEMTN |
| **0C** | SCDSDINIKTGTTDIGSNTTVKTGDLVTYDKENGMHKKVFYSFIDDKNHNKKLLVIRTKGTIAGQYRVYSEEGANKSGLAWPSAFKVQLQLPDNEVAQISDYYPRNSIDTKEYMSTLTYGFNGNVTGDDTGKIGGLIGANVSIGHTLKYVQPDFKTILESPTDKKVGWKVIFNNMVNQNWGPYDRDSWNPVYGNQLFMKTRNGSMKAADNFLDPNKASSLLSSGFSPDFATVITMDRKASKQQTNIDVIYERVRDDYQLHWTSTNWKGTNTKDKWTDRSSERYKIDWEKEEMTN |
| **T117C** | SDSDINIKTGTTDIGSNTTVKTGDLVTYDKENGMHKKVFYSFIDDKNHNKKLLVIRTKGTIAGQYRVYSEEGANKSGLAWPSAFKVQLQLPDNEVAQISDYYPRNSIDTKEYMSTLCYGFNGNVTGDDTGKIGGLIGANVSIGHTLKYVQPDFKTILESPTDKKVGWKVIFNNMVNQNWGPYDRDSWNPVYGNQLFMKTRNGSMKAADNFLDPNKASSLLSSGFSPDFATVITMDRKASKQQTNIDVIYERVRDDYQLHWTSTNWKGTNTKDKWTDRSSERYKIDWEKEEMTN |

**Supplementary Table 2: Sequences and properties of cysteine containing peptides employed for αHL variant modification pre-insertion**

| Modification | Sequence | Properties |
|---|---|---|
| Mod 1 | GSGSGSGSGC | Uncharged, unstructured |
| Mod 2 | DSDGDSDGC | Negatively charged, unstructured |
| Mod 3 | KSKGKSKGC | Positively charged, unstructured |
| Mod 4 | ADARYKSC | Ampholytic, net negatively charged, ATP binding |

**Supplementary Table 3: Sequences of nanoluc reporters used in this study as well as the dark peptide.** $K_D$ and original reporter sequences are taken from Dixon at al. [1] Mean charge calculated at pH 6.5.

| Reporter | Full Sequence | Properties |
|---|---|---|
| r86 | VSGWRLFKKIS | $K_D$ (M) = $0.7x10^{-9}$ |
| r99 | VTGYRLFEKIS | $K_D$ (M) = $1.8x10^{-7}$, +0.91 |
| Dark peptide | VSGWALFKKIS | $K_D$ not measured |
| Unstructured 1 | VTGYRLFEKISGSLGGGGSGGGGSGGGGSAA | $K_D$ not measured, +0.91 |
| Unstructured 2 | VTGYRLFEKISGSLPASPASPASPASPASPA | $K_D$ not measured, +0.91 |
| Helix 1 | VTGYRLFEKISGSGKRQTEREKKKKILAERR | $K_D$ not measured, +6.93 |
| Helix 2 | VTGYRLFEKISGSDKVDEERYDIEAKVTKN | $K_D$ not measured, -1.06 |
| Helix 3 | VTGYRLFEKISGSRRRAAAAAAAAAAAAAAD | $K_D$ not measured, +2.92 |
| Helix 4 | VTGYRLFEKISGSDDDAAAAAAAAAAAAAAR | $K_D$ not measured, -1.09 |

**Supplementary Table 4: Measured currents from MD simulations.**

| Mutation | Modification | Total current (nA) | $K^+$ current (nA) | $Cl^-$ current (nA) |
|---|---|---|---|---|
| WT | - | 0.030 ± 0.002 | 0.009 ± 0.002 | 0.022 ± 0.001 |
| 0C | - | Protonated: 0.027 ± 0.003 | 0.008 ± 0.005 | 0.020 ± 0.006 |
| | | Deprotonated: 0.022 ± 0.004 | 0.011 ± 0.003 | 0.011 ± 0.003 |
| T117C | - | Protonated: 0.025 ± 0.003 | 0.015 ± 0.005 | 0.010 ± 0.008 |
| | | Deprotonated: 0.025 ± 0.002 | 0.015 ± 0.006 | 0.010 ± 0.006 |
| 0C | Mod 2 | 0.016 ± 0.001 | 0.0083 ± 0.0008 | 0.0072 ± 0.0009 |
| | Mod 3 | 0.022 ± 0.002 | 0.0043 ± 0.0009 | 0.017 ± 0.003 |
| | Mod 4 | 0.019 ± 0.002 | 0.0070 ± 0.0008 | 0.012 ± 0.002 |

| | | | | |
|---|---|---|---|---|
| T117C | Mod 2 | 0.0145 ± 0.0008 | 0.010 ± 0.001 | 0.005 ± 0.002 |
| | Mod 3 | 0.015 ± 0.004 | 0.002 ± 0.004 | 0.013 ± 0.001 |
| | Mod 4 | 0.013 ± 0.002 | 0.002 ± 0.001 | 0.011 ± 0.002 |

#### Supplementary figures

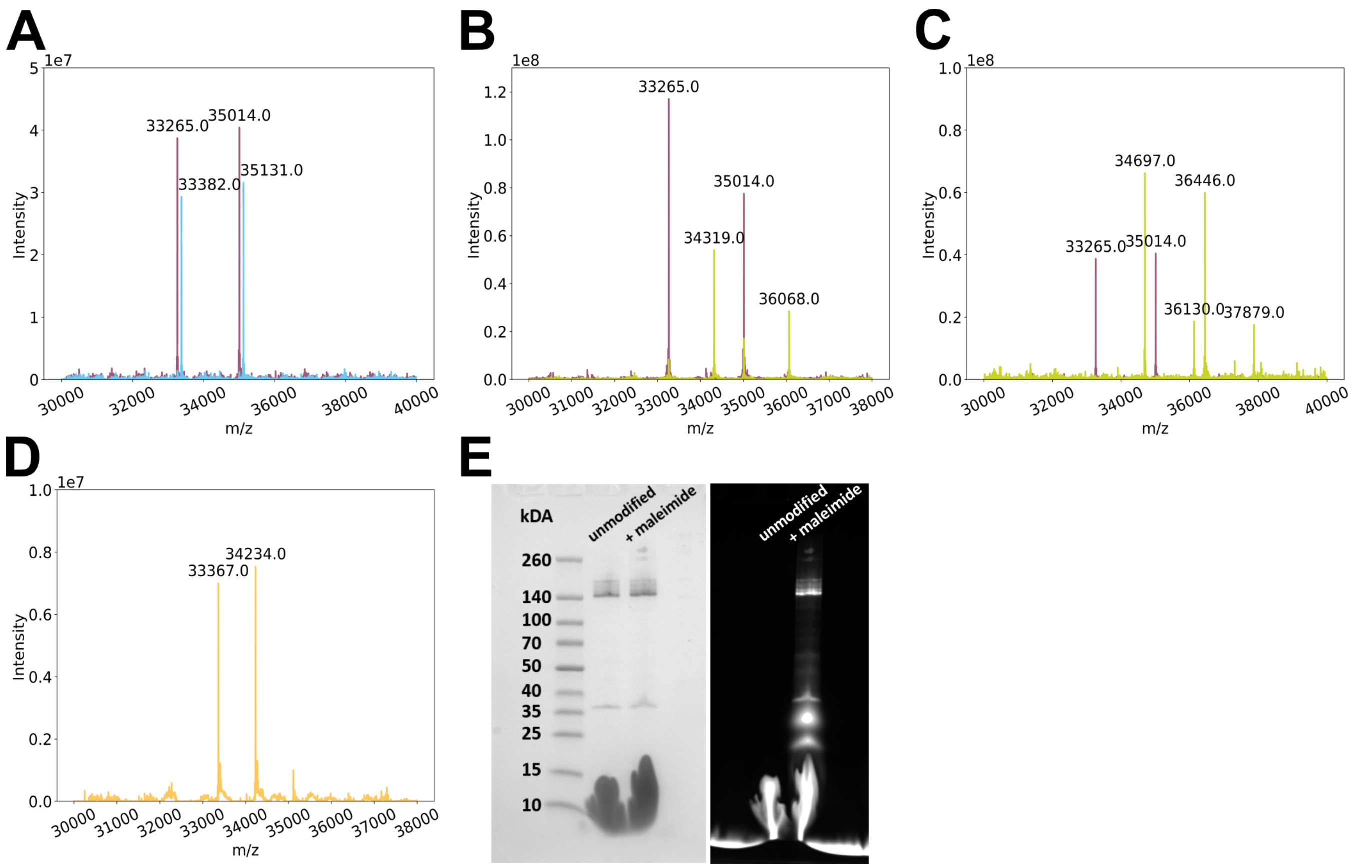


**Supplementary Figure 1: Proof of chemical modification for selected aHL variants and conditions.** Representative intact protein mass spectrometry spectra for A) MTSET modification of aHL T117C monomers, B) Maleimide-Peptide (Mod 4) modification of T117C variant at pH 8, C) maleimide (AQmal) modification of T117C monomers at pH 6.5, D) peptide modification of aHL 0C variant in presence of $CuCl_2$. Here, violet represents the unmodified T177C monomers, green – the monomers incubated with the respective modifying agents and orange the peptide-modified 0C variant. E) SDS page with unmodified and AQmal modified aHL WT, 0C and T117C respectively excited at 488nm and detected at 510nm.

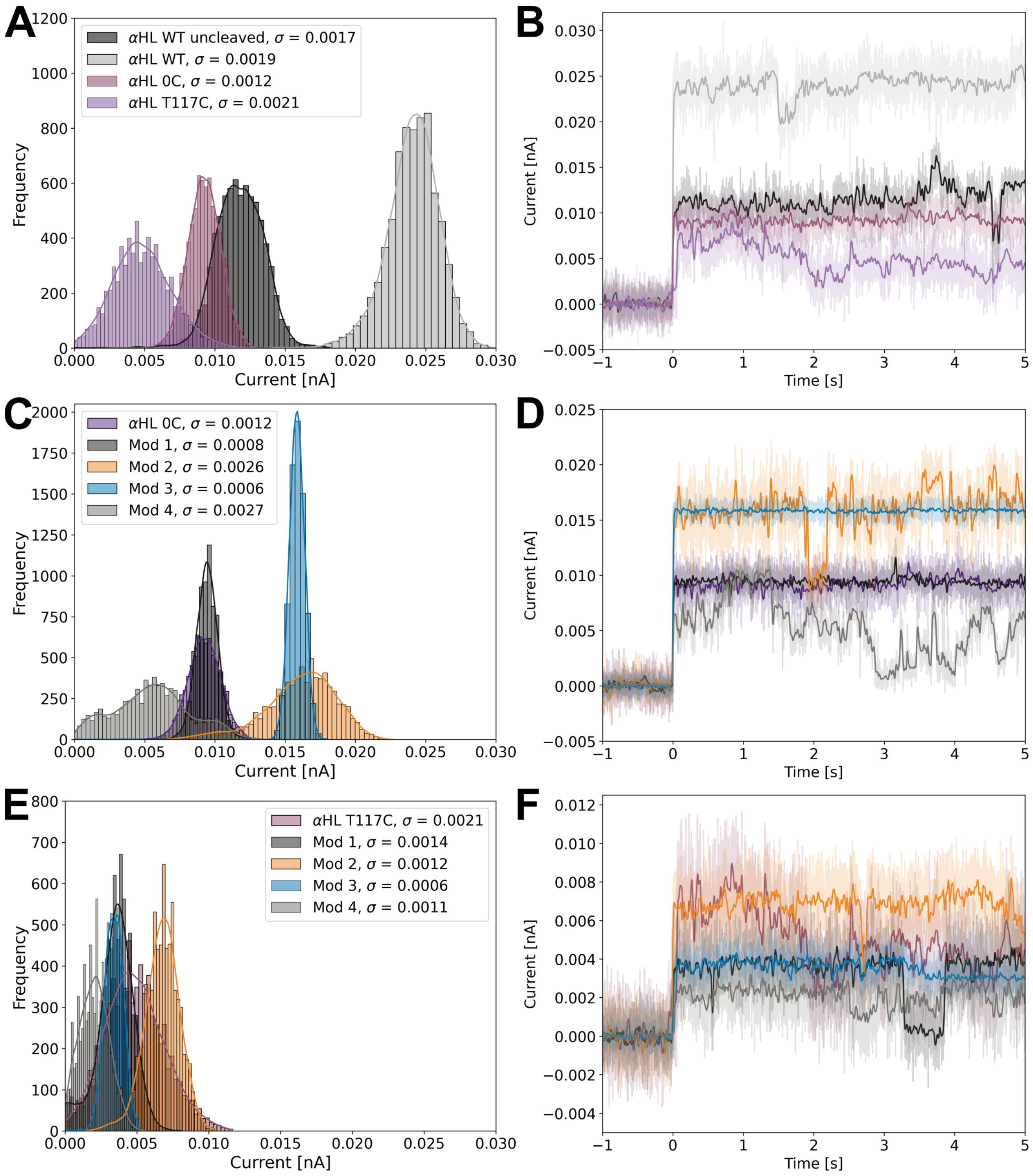


**Supplementary Figure 2: Electrophysiological measurements of single nanopore insertions.** A) Exemplary current distributions recorded for a single insertion of unmodified αHL WT, uncleaved αHL WT as well as its 0C and T117C variants plotted over 6s post insertion. B) The corresponding current traces plotted over 1s pre-insertion and 5s post-insertion. Current histograms (C) and current traces (D) for cysteine peptide modified variants of αHL 0C. Current histograms (E) and current traces (F) for cysteine peptide-modified αHL T117C respectively

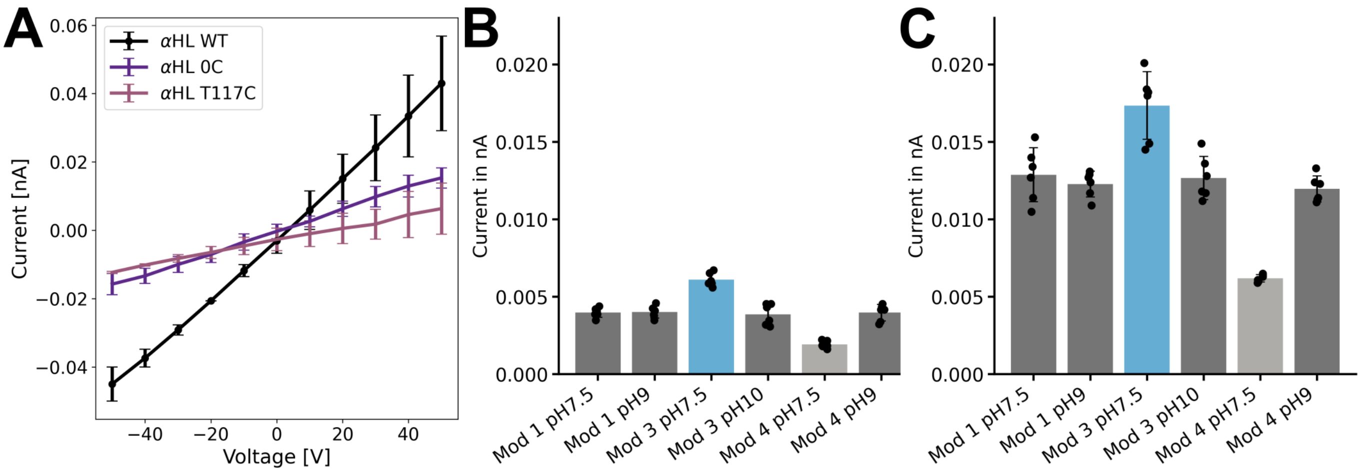


**Supplementary Figure 3: Further electrophysiological characterization of single αHL insertions**. A) IV curves measured for n=3 independent insertions of the studied αHL variants. B) Changes in mean current for n=6 independent insertions of αHL T117C variant functionalized with different cysteine containing peptides. C) Respective changes for αHL 0C variant. Here, dark grey is used to show net 0 charge, blue net positive charge and light grey is used to denote ampholythic peptides.

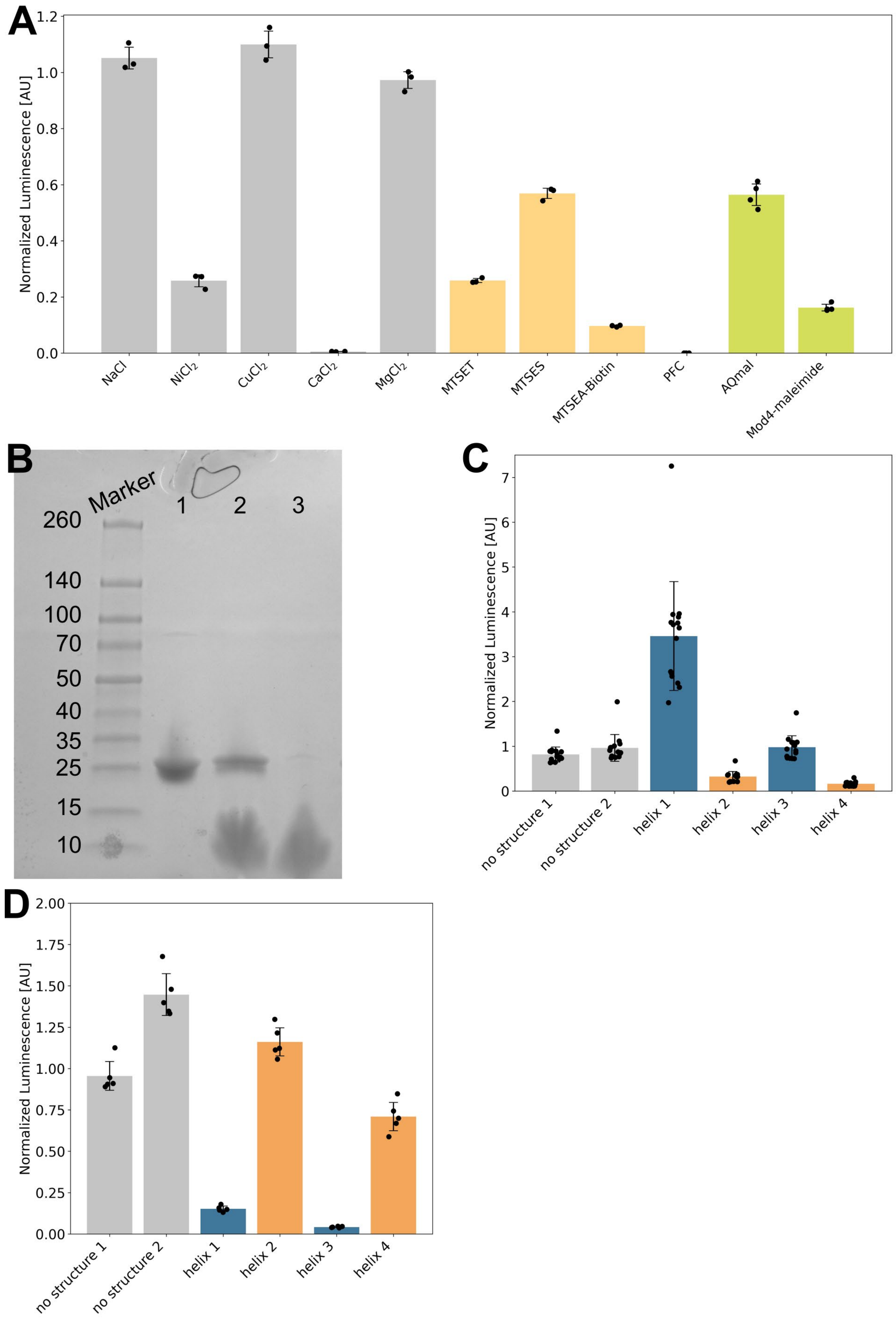
A
Normalized Luminescence [AU]
0.0
0.2
0.4
0.6
0.8
1.0
1.2
NaCl
NiCl2
CuCl2
CaCl2
MgCl2
MTSET
MTSES
MTSEA-Biotin
PFC
AQmal
Mod4-maleimide
B
Marker
1
2
3
260
140
100
70
50
40
35
25
15
10
C
Normalized Luminescence [AU]
0
1
2
3
4
5
6
7
no structure 1
no structure 2
helix 1
helix 2
helix 3
helix 4
D
Normalized Luminescence [AU]
0.00
0.25
0.50
0.75
1.00
1.25
1.50
1.75
2.00
no structure 1
no structure 2
helix 1
helix 2
helix 3
helix 4

**Supplementary Figure 4: Characterization of the nanoluc system and engineered reporters in bulk luminescence assays.** A) effect metal salts, as well as modifying reagents have on the luminescence output of the system. Here, all metal data is plotted in grey, the MTS reagent data is plotted in orange and maleimide functionalization in green. All data is normalized to the r99 output in absence of additional reagents. The assay was performed in n=3 technical replicas and plotted for t=15min. B) non-reducing SDS-PAGE, with line 1 representing the unencapsulated LgBit (100%), line 2 – DOPC:DOPG:DOPE vesicles with LgBit encapsulated, line 3 DOPC vesicles with LgBit encapsulated. C) luminescence output of the engineered reporter variants normalized to r99 output. D) split nanoluciferase bulk assay luminescence output of engineered reporter peptides in presence of DOPC:DOPG:DOPE (40:30:30 molar ratio) LUVs. The data is plotted for t=30min with n=3 independent assays with n=5 technical replicas each pooled.

**Supplementary Figure 5: Estimation of the ionic current from the slope of a linear fit to the cumulative curve as a function of time.** Data are shown for the mutation 0C, without peptide, with all cysteines protonated. Cumulative currents are shown for potassium (blue) and chloride (orange), with the corresponding linear fits shown as dark dashed lines. The estimated average currents were 0.008 +/0.005 nA for potassium, and 0.022 +/ 0.006 nA for chloride, with errors computed from 200 ns blocks.

#### Supplementary Text 1: Ion diffusion coefficients and effects on flux

The diffusion coefficients of potassium and chloride ions in water were computed from molecular dynamics simulations performed at a KCl concentration of 3 M. The systems were built using CHARMM-GUI [2] solution builder, generating cubic boxes of side lengths 5, 6, 7 and 8 nm, and the CHARMM36m force field [3] was used. Simulations were performed with GROMACS 2024.0 [4]. Each system was first energy-minimized using the steepest-descent algorithm, followed by 125 ps of NVT equilibration using a 1 fs time step at 298.15 K with the v-rescale thermostat [5] and a coupling time constant of 1.0 ps. A 20 ps NPT equilibration using 2 fs time step was then performed at 298.15 K and 1 bar using the C-rescale barostat [6], with a coupling time constant of 5.0 ps. Production simulations were carried out in the NVT ensemble at 298.15 K, using a 2 fs time step, for 200 ns for the 5 nm box, 100 ns for the 6 nm box, and 50 ns for the 7 and 8 nm boxes.

Mean-square displacements were calculated separately for potassium and chloride ions, using the gmx msd command, averaging over all ions of the same species in each system. Diffusion coefficients were then obtained for each box size and extrapolated to infinite box size following the Yeh and Hummer approach [7]. The diffusion coefficient obtained under periodic boundary conditions was assumed to vary linearly with the inverse box length $1/l$. The diffusion coefficient at infinite box size was thus obtained from the intercept of a linear fit of $D(L)$ as a function of $1/l$ at $1/l = 0$, allowing comparison with experimental diffusion coefficients [8,9].

Ion self-diffusion coefficients computed for the 3 M KCl solution at different box sizes allowed us to extrapolate to infinite system size and remove the effects of periodic boundary conditions. The simulations were performed at higher concentration to ensure faster and better sampling. Computed diffusion coefficients were $(1.59 \pm 0.12) \cdot 10^{-5}$ cm$^2$/s and $(1.64 \pm 0.09) \cdot 10^{-5}$ cm$^2$/s for $K^+$ and $Cl^-$ in a 5 nm box, $(1.74 \pm 0.01) \cdot 10^{-5}$ cm$^2$/s and $(1.82 \pm 0.25) \cdot 10^{-5}$ cm$^2$/s for $K^+$ and $Cl^-$ in a 6 nm box, $(1.613 \pm 0.008) \cdot 10^{-5}$ cm$^2$/s and $(1.80 \pm 0.10) \cdot 10^{-5}$ cm$^2$/s for $K^+$ and $Cl^-$ in a 7 nm box, and $(1.73 \pm 0.04) \cdot 10^{-5}$ cm$^2$/s and $(1.71 \pm 0.06) \cdot 10^{-5}$ cm$^2$/s for $K^+$ and $Cl^-$ in a 5 nm box. Considering that $D \sim 1/l$, where $l$ is the side length of a simulation box, we obtained a diffusion coefficient at infinite box size by calculating the intercept of linear regression of data points [7]. Estimated values of ion self-diffusion coefficients in 3 M KCl aqueous solution are $1.86 \cdot 10^{-5}$ cm$^2$/s for $K^+$, and $1.93 \cdot 10^{-5}$ cm$^2$/s for $Cl^-$ ions. Compared to experimental values of self-diffusion coefficients in 3 M KCl [8,9] of $1.84 \cdot 10^{-5}$ cm$^2$/s for $K^+$ and $1.87 \cdot 10^{-5}$ cm$^2$/s for $Cl^-$, we conclude that the diffusion coefficients, and sequentially computed ion currents, are overestimated by 1 and 3%, respectively
